# Single Cell Transcriptomics Reveals Dysregulated Cellular and Molecular Networks in a Fragile X Syndrome model

**DOI:** 10.1101/2020.02.12.946780

**Authors:** Elisa Donnard, Huan Shu, Manuel Garber

## Abstract

Despite advances in understanding the pathophysiology of Fragile X syndrome (FXS), its molecular bases are still poorly understood. Whole brain tissue expression profiles have proved surprisingly uninformative. We applied single cell RNA sequencing to profile a FXS mouse model. We found that FXS results in a highly cell type specific effect and it is strongest among different neuronal types. We detected a downregulation of mRNAs bound by FMRP and this effect is prominent in neurons. Metabolic pathways including translation are significantly upregulated across all cell types with the notable exception of excitatory neurons. These effects point to a potential difference in the activity of mTOR pathways, and together with other dysregulated pathways suggest an excitatory-inhibitory imbalance in the FXS cortex which is exacerbated by astrocytes. Our data demonstrate the cell-type specific complexity of FXS and provide a resource for interrogating the biological basis of this disorder.

## Introduction

Fragile X syndrome (FXS) is the most common inherited form of intellectual disability and autism spectrum disorder (ASD). The disease results from the silencing of a single gene (*Fmr1*), which encodes the RNA-binding protein FMRP ^1^. The loss of FMRP leads to a neurodevelopmental disorder with an array of well characterized behaviour and cellular abnormalities, such as impaired cognitive functions, repetitive behaviours, altered synaptic morphology and function ^2^; many of which are reproduced in *Fmr1*-KO mouse models ^3^.

The molecular pathophysiology of FXS and FMRP function has been the subject of numerous studies over the past decades ^4,5^. The most extensively studied function of FMRP is its role as a translational repressor. FMRP is critical to hippocampal long-term synaptic and spine morphological plasticity, dependent on protein synthesis. More specifically, the absence of FMRP leads to an exaggerated long-term synaptic depression, induced by the metabotropic glutamate receptor 5 (mGLUR5-LTD) ^6^. However, several ambitious clinical trials that aimed to suppress translation or inhibit mGluR pathways have thus far failed ^7^. The significance of FMRP’s role as a translation repressor at synapses is not without challenges. First, only a few mRNAs that are bound by FMRP showed a consistent increase in protein levels upon loss of FMRP, and increased levels of proteins are not always pathogenic ^8^. More importantly, focusing on FMRP’s translational function in dendritic synapses overlooks the fact that the great majority of this protein is located in the cell soma ^9^. Indeed, a wide range of research has associated FMRP to multiple steps of the mRNA life cycle, including pre-mRNA splicing ^10^, mRNA editing ^11,12^, miRNA activity ^13,14^, and mRNA stability ^15,16^. Additionally, FMRP may function outside the RNA-binding scope, by chromatin binding and regulating genome stability ^17,18^, as well as directly binding to and regulating ion channels ^19,20^.

Most of the above-mentioned studies focus on FMRP’s function in neurons, and rightly so, as neurons have the highest FMRP protein levels in the brain ^9^. Evidence from clinical studies with FXS patients and from mouse models of the disease supports the view that neurons are the main affected cell type ^21,22^. However, to add to the complexity, this is not to say that the other cell types in the brain do not express FMRP or are not affected upon loss of it. Indeed, astrocytes, oligodendrocyte precursor cells, and microglia express FMRP in a brain structure and development-dependent manner ^23^. FMRP-depleted astrocytes are more reactive ^24^, and their deficits alone may account for some of the phenotypes seen in Fragile X neurons particularly during development ^25–28^.

This profound body of work over decades has collectively pieced together a complex picture for FMRP’s functions. However, this also poses a challenge to building an overview of the molecular impact of Fragile X. Here we present our effort to bridge this gap using an unbiased approach to survey the Fragile X brain. We took advantage of the power of single cell RNA-seq to determine which cells are affected at an early postnatal development stage, using the transcriptome as a sensitive reflection of the cellular status.

We profiled the transcriptome of over 18,000 cells from the cerebral cortex of wild type (WT) and *Fmr1*-knock out mice (*Fmr1*-KO) at postnatal day 5. Our findings present new insights into the cell type specific consequences of FXS. We detected a heterogeneity in the response of different cell types to the loss of FMRP. We find in particular a higher impact on the expression of mRNAs previously identified as FMRP binding targets in the brain (Darnell et al 2011), and we show that this effect is prominent in neurons compared to other cells. We detect a divergent response of pathways downstream of mTOR signaling across different neuron subtypes, which suggests that excitatory neurons do not display a hyperactivation of this pathway. Taken together with the observed dysregulation of synaptic genes in astrocytes as well as neurons, our results suggest an impact in cell-cell communication that can result in a cortical environment of greater excitability.

## Cell type proportions are not impacted in the neonatal *Fmr1*-KO cortex

To capture early molecular events in the Fragile X syndrome, we performed single cell RNA-seq using the InDrop system ^29^, from the cortex of both *Fmr1*-knock out (*Fmr1*-KO) and wild type (WT) FVB animals at postnatal day 5 (P5, Figure 1a, Methods), a critical period in cortical development for neuronal and synaptic maturation ^30–32^. After stringent filtering, we obtained 18,393 cells for which we detected an average of 1,778 genes and 3,988 transcripts (see Methods). After unbiased clustering we classified these cells into seven major cell types, with specific expression of established markers^33–36^ (Figure S1a; Methods).

**Figure 1.**
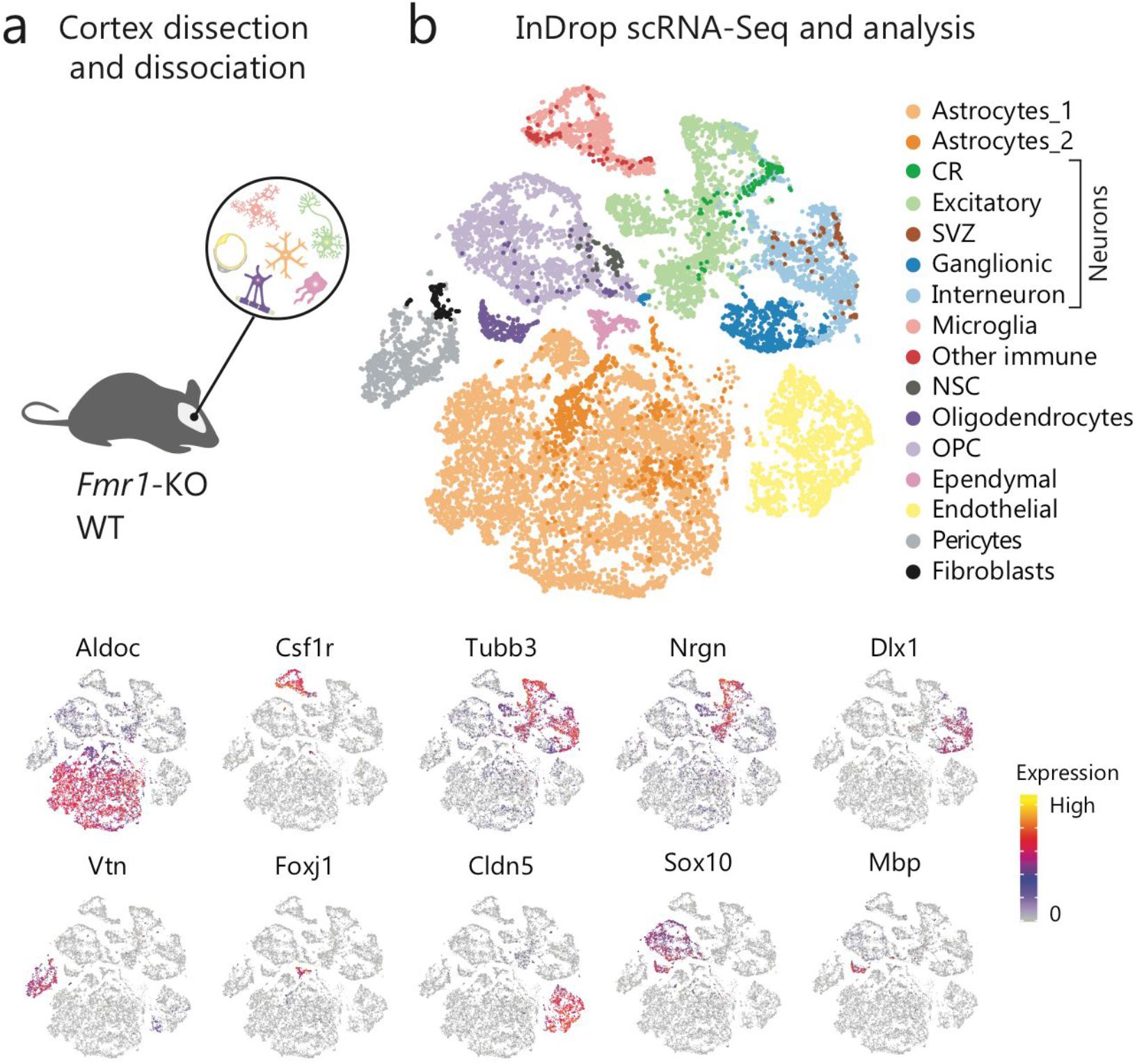
Cortical cell types in *Fmr1*-KO and WT mice. **a)** Overview of the sample collection **b)** tSNE representation of cells colored by cell type (top) and colored by expression level of example maker genes (bottom).

We did not detect a genotype bias in the clustering (Figure S1b). Additionally, all major cell types were detected in both *Fmr1*-KO and WT animals, with consistent proportions across individual mice (Figure S1c), indicating that *Fmr1*-KO mice do not have large scale cell differentiation deficits at this stage, consistent with previous reports ^9^. To better dissect the effect of FMRP loss on different cell types, we independently re-clustered the broad cell types. We identified a total of 18 distinct populations (Figure 1b, S1d-g; Methods). These populations include vascular cells (endothelial cells and pericytes), fibroblasts, ependymal cells, different neuron subtypes and glial cells (Figure 1b). Neuron clusters were classified as excitatory neurons (Figure S1d, Excitatory, *Nrgn*^*+*^ *Lpl*^*+*^ *Slc17a6*^*+*^), immature interneurons (Figure S1d, Interneuron: *Gad1*^*+*^ *Dlx1*^*+*^ *Htr3a*^*+*^), ganglionic eminence inhibitory precursors (Figure S1d, Ganglionic: *Ascl1*^*+*^ *Dlx2*^*+*^ *Top2a*^*+*^), Cajal Retzius cells (Figure S1d, CR: *Reln*^*+*^ *Calb2*^*+*^ *Lhx5*^*+*^) and subventricular zone migrating neurons (Figure S1d, SVZ: *Eomes*^*+*^ *Sema3c*^*+*^ *Neurod1*^*+*^). We detected two immature astrocyte populations, including a small group with markers of reactive astrocytes (Figure S1e, Astrocytes_1: *Aqp4*^*+*^ *Aldoc*^*+*^ *Apoe*^*+*^; Astrocytes_2: *Ptx3*^*+*^ *Igfbp5*^*+*^ *C4b*^*+*^). Immune cells detected consist largely of microglia (Figure S1f, Microglia: *Cx3cr1*^*+*^ *P2ry12*^*+*^ *Csf1r*^*+*^), but we also identified small clusters of T cells (Figure S1f, T Cells: *Cd3d*^*+*^ *Ccr7*^*+*^ *Cd28*^*+*^), border macrophages (Figure S1f, Border Macrophages: *Lyz2*^*+*^ *Msr1*^*+*^ *Fcgr4*^*+*^) and a population of microglia-like cells marked by high expression of Ms4a cluster genes (Figure S1f, Microglia Ms4a+: *Ms4a7*^*+*^ *Ms46b*^*+*^ *Mrc1*^*+*^). The initial clusters containing oligodendrocytes were classified as mature oligodendrocytes (Figure S1g, Oligodendrocytes: *Mbp*^*+*^ *Sirt2*^*+*^ *Plp1*^*+*^), oligodendrocyte progenitor cells (Figure S1g, OPC: *Pdgfra*^*+*^ *Olig2*^*+*^ *Sox10*^*+*^) and a small group of neural stem cells (NSC: *Btg2*^*+*^ *Dll1*^*+*^ *Dbx2*^*+*^).

## Loss of FMRP results in small expression changes across a broad spectrum of processes

We examined differential expression between *Fmr1*-KO and WT mice for each cell type identified. In total, we identified 1470 differentially expressed (DE) genes (FDR < 0.01 and fold change >= 1.15x) in one or more cell types (Figure S2a-b; Table S1). The majority of DE genes showed small fold changes (mean fold change = 1.3x). The effect of FMRP loss seems drastically different across cell types as most differentially expressed genes were found in neurons and endothelial cells (Figure S2a). This result does not seem to be dependent on statistical power, given that the number of DE genes detected did not correlate to the number of cells available for different cell types.

Given the broad but low effect size resulting from loss of FMRP, we focused on quantifying the impact on annotated pathways. To this end, we performed a gene set enrichment analysis (GSEA; see Methods). Consistent with the larger effect observed using differential gene expression analysis, most significantly dysregulated processes are also found in neurons, further suggesting that neurons are the cells most impacted by FMRP loss (Figure 2a-b, Figure S2c-d, Table S2).

**Figure 2.**
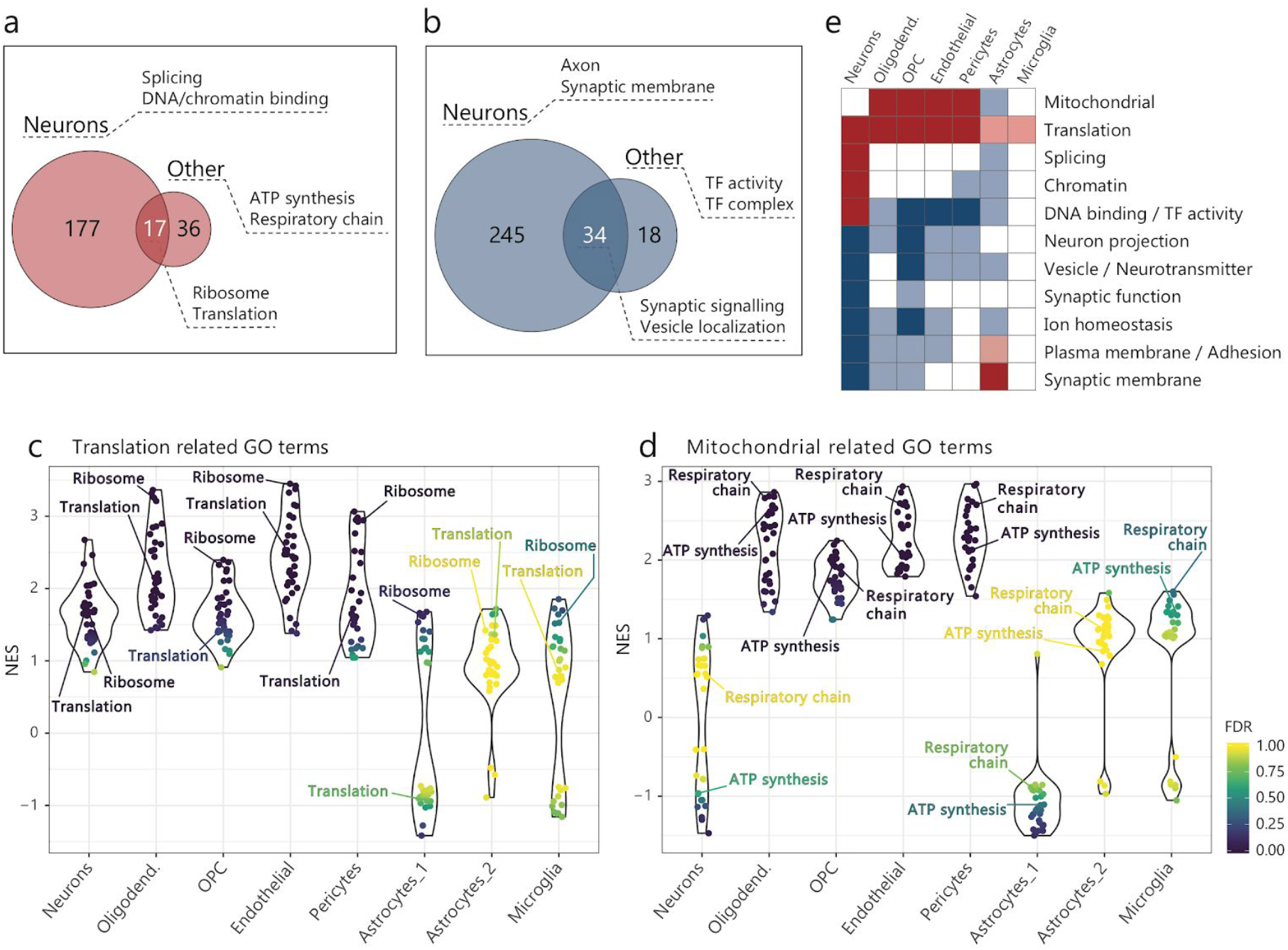
Misregulated Gene Ontology categories per cell type reveal higher impact in *Fmr1*-KO neurons. **a-b)** Comparative diagram of upregulated (a) or downregulated (b) GO categories in neurons and other cell types. Keywords and terms from the top enriched GO categories for each ontology are shown. **c-d)** Comparison of normalized enrichment scores (NES) across cell types for GO terms involved in translational (c: e.g. Ribosome, GO:0005840; Translation, GO:0006412) or mitochondrial (d: e.g. Respiratory chain, GO:0070469; ATP synthesis = ATP synthesis coupled electron transport, GO:0042773) processes. **e)** Summary diagram of enriched processes (dark red/blue; FDR<0.01) and trends (light red = NES>1, FDR<0.3; light blue = NES<−1, FDR<0.3) in other cell types.

We find that metabolic processes are upregulated in most FMRP deficient cell types (e.g. Translation, Ribosome, Respiratory Chain Complex and Oxidative Phosphorylation). Translational processes are significantly upregulated in neurons, vascular cells (endothelial and pericytes), oligodendrocytes and OPCs. However, upregulation is clear in most cell types even when the difference did not reach a significant level (Figure 2c). Similarly, mitochondrial pathways are some of the most strongly upregulated processes in multiple cell types including OPCs, oligodendrocytes, endothelial cells and pericytes (Figure 2d). Transcriptional activation of genes involved in ribosome biogenesis ^37^ as well as oxidative phosphorylation ^38^ are known downstream effects of the mTOR pathway. This upregulation across multiple cortical cell types is in agreement with previous reports of elevated phosphorylation of mTOR in *Fmr1*-KO mice ^39,40^, and subsequent increase in signalling through the mTOR complex 1 pathway. The increased abundance of mRNAs for translation related factors in all cortical cell types also agrees with the well established impact of FMRP on protein synthesis ^39,41^.

On the other hand, there is a common trend of downregulation across many cell types for genes involved in synaptic signaling, vesicle transport and ion homeostasis (Figure 2e). Most of these processes are significantly downregulated in neurons and OPCs, and include common downregulated genes (e.g. Apoe, Sar1b, Clstn1, Rab5a and Sqstm1 involved in vesicle pathways), suggesting that intracellular transport of molecules is impaired in multiple cell types. Synaptic signaling and cell surface proteins notably respond differently in astrocytes, and are discussed below.

Taken together, this reveals a transcriptional landscape in the neonatal *Fmr1*-KO cortex where not all cell types are affected equally. Most genes are differentially expressed in a cell type specific manner, however they are often involved in related processes across cell types. There is excessive expression of factors involved in biomolecule production (protein and ATP synthesis) in multiple cell types. Conversely, there is insufficient expression of factors important for developing cellular communication and maintaining ion homeostasis in multiple cell types, particularly in neurons (Figure 2e). These related processes that are impacted in multiple cell types in the *Fmr1*-KO cortex collectively point to enhanced growth and metabolism while suggesting hindered or delayed network development through impaired cell-cell communication.

## Multiple pathways display opposite effects in astrocytes and neurons and suggest hyperexcitability

We detect many processes where neurons and astrocytes show opposite effects, revealing a surprisingly different impact of the loss of FMRP in these two cell types. Genes encoding for synaptic signaling and cell surface proteins are in general downregulated in neurons and other cell types, while upregulated in astrocytes (Figure 3a). The main dysregulated genes in these processes include a large number of known receptor-ligand pairs involved in cross-cell signaling and their dysregulation could contribute to an imbalanced extracellular neuro and gliotransmitter homeostasis (Figures 3b-c; leading edge analysis, GSEA). Upregulated genes in astrocytes contributing to this enrichment include ephrin and GABA receptors as well as other solute carriers. Related genes, however, are downregulated in neurons. For example, we detected a downregulation of Efna5 and Epha7 in *Fmr1*-KO neurons, both critical to cortex development and plasticity and involved in the establishment of corticothalamic projections ^42–44^. This downregulation could therefore be involved in the deficit of cortical-subcortical circuits reported in individuals with FXS ^45^. Conversely, we observed an upregulation of Ephb3 and Epha4 receptors in astrocytes, which are activated by ephrin-B produced by neurons (Figure 3b). Activation of these receptors leads to enhanced release of D-serine (NMDA receptor coagonist) and glutamine (uptaken by excitatory terminals and converted to glutamate)^46,47^. Similarly, we detected a downregulation of Slc1a4 (Asct1) in neurons, which could also contribute to reduced clearance of D-serine ^48^.

**Figure 3.**
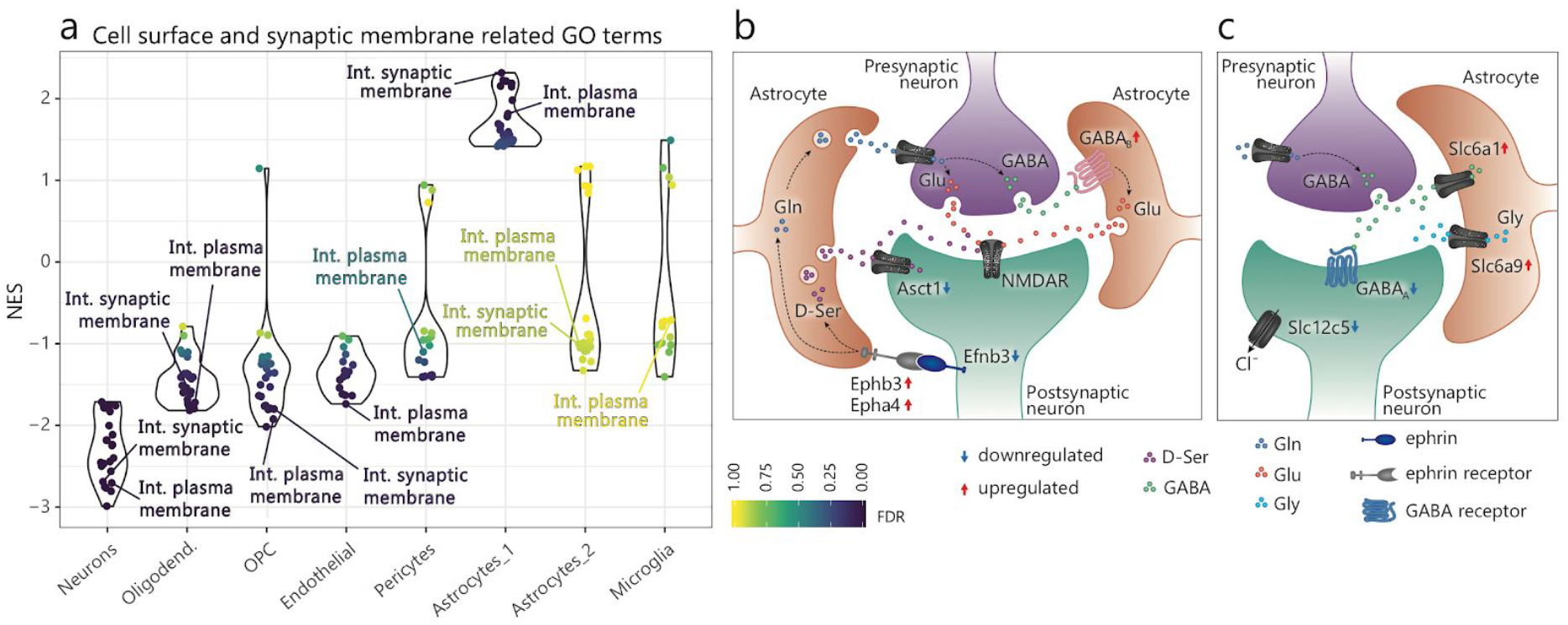
Dysregulated genes in *Fmr1*-KO astrocytes and neurons involved in synaptic functions. **a)** Comparison of normalized enrichment scores (NES) across cell types for all GO terms that show opposite signals in astrocytes and neurons (e.g. Int. synaptic membrane = Integral component of synaptic membrane, GO:0099699; Int. plasma membrane = Intrinsic component of plasma membrane, GO:0031226). **b-c)** Summary diagram of strongest dysregulated genes in astrocytes and neurons involved in synaptic signaling and regulation which suggest an increased release of excitatory (b) and a higher clearance of inhibitory (c) glio and neurotransmitters in the synaptic environment. D-Ser = D-Serine; GABA = Gamma aminobutyric acid; Gln = Glutamine; Glu = Glutamate; Gly = Glycine.

Further, we observed an upregulation of Gabbr1 and Gabbr2 in *Fmr1*-KO astrocytes, suggesting enhanced glutamate release resulting from the activation of GABA_B_ receptors (Figure 3b) ^49^. Other genes that could lead to enhanced clearance of GABA and glycine (both inhibitory neurotransmitters) from the neuronal environment are also upregulated in astrocytes, such as Slc6a1 (GAT-1), and Slc6a9 (GlyT-1) (Figure 3c). Our data also suggest a reduced sensitivity to environmental GABA inhibitory signals in neurons, which show downregulation of a GABA_A_ receptor gamma subunit (Gabrg2). Previously, in both fly and mouse models of FX, multiple subunits (including gamma) were reported to have reduced expression ^50^. We additionally detected reduced expression of Slc12c5 (KCC2) in *Fmr1*-KO neurons, a transporter crucial for the switch of GABA from being excitatory to inhibitory ^51^, which could underlie the delayed switch in GABA polarity seen in *Fmr1*-KO mice ^52^. Collectively, these changes point to an imbalanced extra and intra-neuronal environment that favors excitation over inhibition (Figures 3b-c). Indeed, cortical hyperexcitability is believed to be the biological basis of ASD and epilepsy, including FXS^53^.

Several other observed examples of dysregulated solute carriers in multiple cell types may contribute to the FXS phenotypes of mRNA translation defects and autistic behaviors through pathways other than regulation the glio and neurotransmitters as discussed above. These include genes downregulated in oligodendrocytes, such as Slc1a2 (GLT-1), involved in glutamate regulation and critical for white matter development ^54,55^, or downregulated in endothelial cells, such as Slc7a5 (LAT-1), a mediator of amino acid uptake which can impact the amino acid profile and mRNA translation in the brain, causing neurological abnormalities ^56^. Astrocytes in turn, show an increased expression of Slc30a10 (ZnT10), responsible for Zn^2+^ and Mn^2+^ transport to the extracellular space ^57^. High cellular levels of Zn^2+^ and Mn^2+^ in turn induce the transcription of metallothioneins ^58^. We observed upregulation of Slc30a10 in astrocytes, which would predict downregulation of metallothioneins. Indeed, Mt1 and Mt2 are among the most strongly downregulated genes in astrocytes (Figure S2b). Metallothioneins play a role in the protection of the central nervous system in response to injury, and neuronal recovery is impaired in their absence ^59^.

Our data points to a picture in which cell types are impacted widely in different and sometimes compensatory ways, and these expression changes are moderate. This may explain why previous whole brain and whole cortex assays have provided underwhelming insights. The ability to have a cell type specific expression profile highlights the wide range of the effects of FMRP loss that result in strong pathology.

## FMRP loss impacts neuronal homeostasis

A total of 473 categories (88%) were affected in neurons (Figure 2a-b, S2c-d). We found that the main neuron-specific upregulated processes correspond to RNA splicing, DNA binding and chromatin organization (Figure 4a). Upregulated genes in RNA splicing categories encode core spliceosomal proteins ^60^, including RBM39, Snrnp200, Hnrnpn, Srsf3 and Srsf4. Upregulation of these factors has been previously linked to an increased rate of proliferation ^61–63^. Similarly, the most strongly upregulated genes associated with DNA binding, chromatin and other significant transcriptional regulation terms include many regulators of proliferation (Hmgb2, Top2a, Mcm7, Pcna) and of neuron differentiation (Insm1, Dlx2, Tead1, Tshz1, Tcf12) ^64–66^.

**Figure 4.**
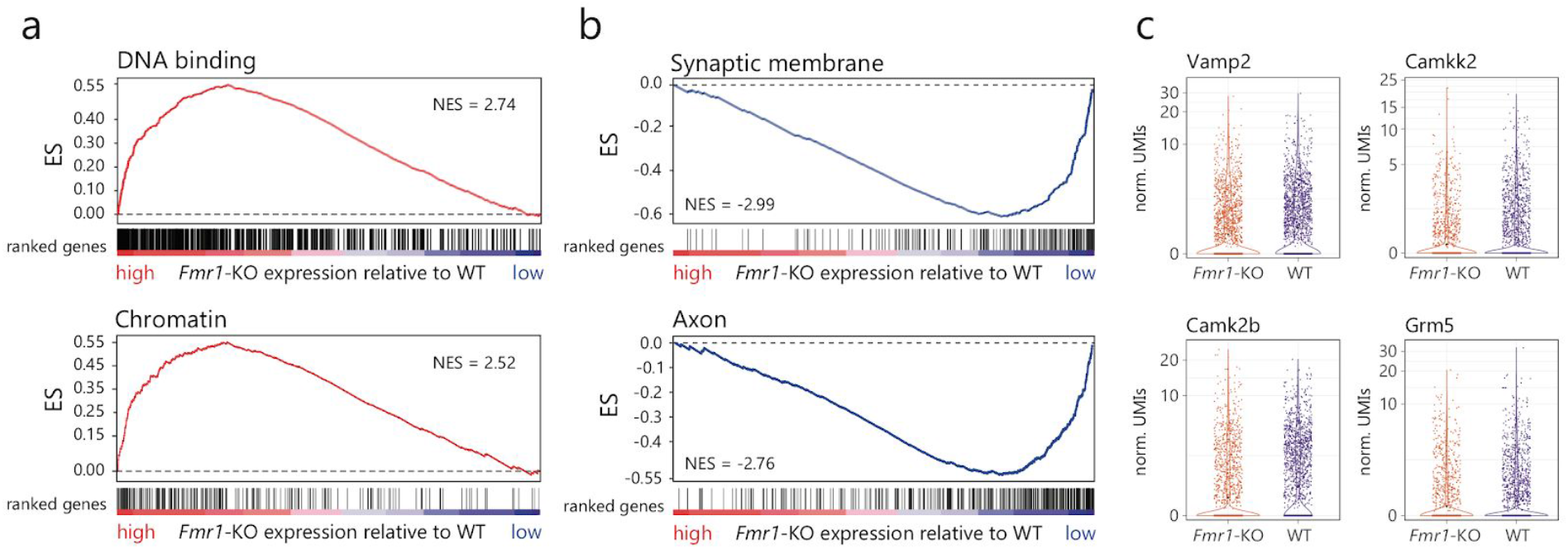
FMRP loss impacts neuronal homeostasis. **a)** Example neuron-specific upregulated gene groups: DNA binding (GO: 0003677; top), and chromatin (GO:0000785; bottom). **b)** Example neuron-specific downregulated gene groups: synaptic membrane (GO:0097060; top) and axon (GO:0030424; bottom). ES = GSEA Enrichment Score. NES = GSEA Normalized Enrichment Score. **c)** Expression in all neuronal cells for example genes downregulated in *Fmr1*-KO involved in synaptic and neuron projection processes.

More categories are specifically downregulated in neurons (n=245, 51%), which include functions related to synaptic and membrane components, ion channels and neuron projections (Figure 4b). This broad impact in synaptic processes is consistent with the idea that FXS is a neuronal disorder that results from a suboptimal balance in connectivity ^67^. The top downregulated genes related to synapses and neuronal projections (Figure 4c) include Vamp2, a key component of synaptic vesicle trafficking ^68^; Camkk2 and Camk2b, which are both involved in the formation of dendritic spines ^69^; and Grm5, which encodes for the mGluR5 receptor, and noticeably is one of the main proposed therapeutic targets for FXS ^70^. Reduced abundance of these and other synaptic and neuronal projection related mRNAs reveals multi-leveled transcriptome deficits (vesicle transport, cell morphology and receptors, affecting both pre and postsynaptic structures), which could be behind the known deficits of FXS neurites and synaptic development ^4^.

## FMRP bound mRNAs are downregulated in *Fmr1*-KO neurons

FMRP binds directly hundreds of mRNAs identified in the mouse brain^71–73^ or in cell lines ^74,75^. The current model suggests that FMRP binding leads to translational repression of its bound mRNAs (Darnell 2011, Chen et al 2014), although this repression was only validated for a handful of direct targets ^8^. We focused on the impact of FMRP loss on mRNA abundance of its direct binding targets. Out of the 842 mRNAs bound by FMRP in the mouse brain ^72^, we detected 839 expressed by at least one cell type in our data. We compared the fold change between *Fmr1*-KO and WT for FMRP bound mRNAs to all other expressed. Consistent with the strong effect of FMRP loss in neurons, FMRP bound mRNAs show a stronger downregulation in neurons compared to other expressed mRNAs (p < 10^−57^, Wilcoxon rank-sum), while for other cell types there is little difference (Figure 5a, S3a).

**Figure 5.**
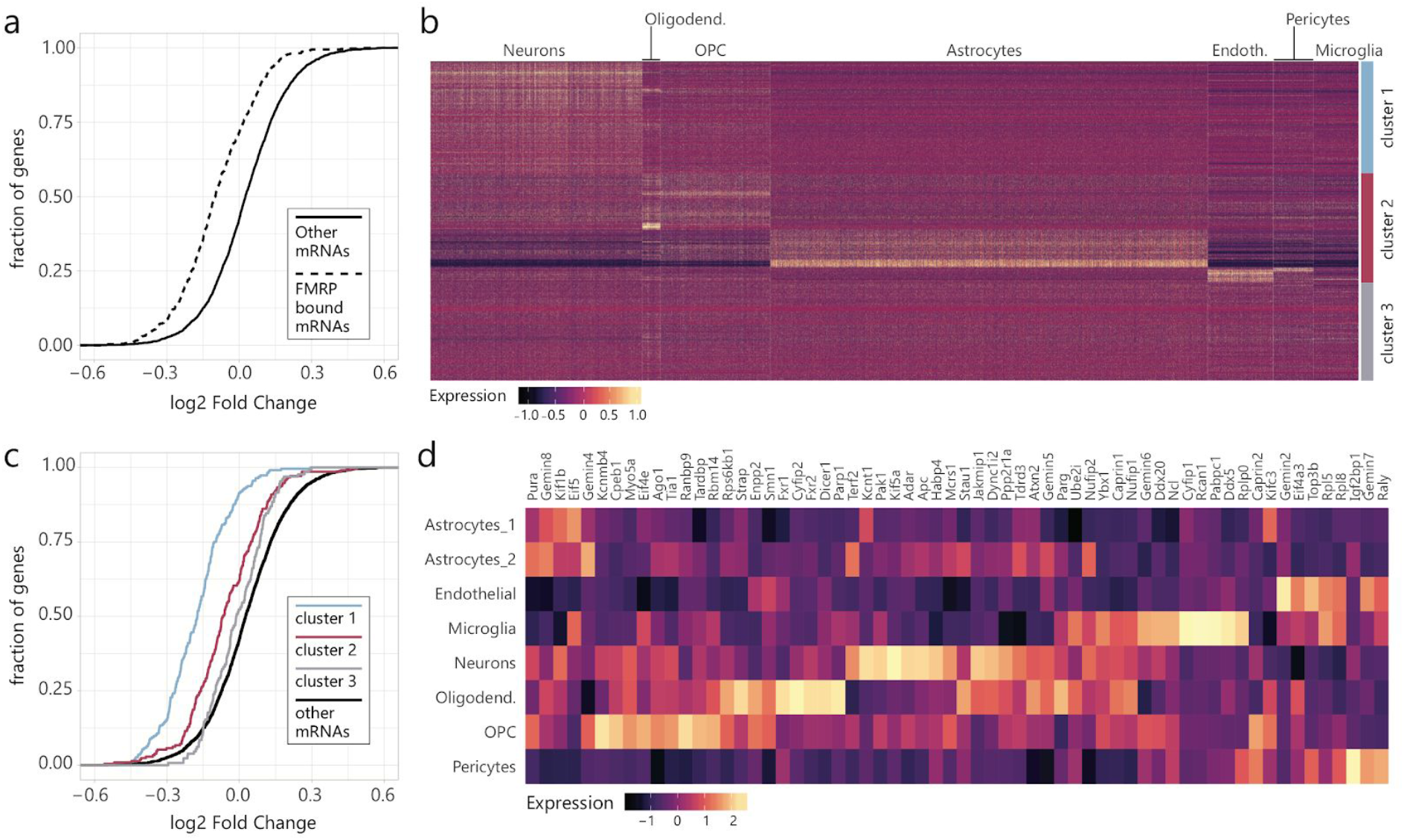
FMRP bound mRNAs show greater downregulation in neurons. **a)** Cumulative distribution plot of the log2 fold change between *Fmr1*-KO and WT gene expression in neurons. Dashed lines represent mRNAs annotated as FMRP bound by Darnell et al. expressed in that cell type. Full lines represent all other expressed genes in that cell type. **b)** Heatmap of expressed FMRP bound mRNAs in WT cortex cells. Genes were clustered based on their expression pattern across cell types and resulting clusters comprise genes expressed at higher levels in neurons (cluster 1), non-neuronal cell types (cluster 2) and without cell type specificity (cluster 3). **c)** Cumulative distribution plot of the log2 fold change between *Fmr1*-KO and WT gene expression in neurons. Colored lines represent each specific subset of expressed genes: Neuron FMRP bound mRNAs, which show highest expression in neurons (blue, cluster 1, Figure 5b); Non-neuronal FMRP bound mRNAs, which show highest expression in other cell types (pink, cluster 2, Figure 5b); Non-specific FMRP bound mRNAs (grey, cluster 3, Figure 5b), which are expressed at similar levels by all cortical cell types; and all other mRNAs expressed in neurons (black). **d)** Expression levels of known protein binding partners of FMRP across cell types. Aggregated bulk values were calculated per cell type (UMIs per million).

This steeper downregulation of FMRP bound mRNAs in neurons could result from higher expression levels of these mRNAs in neurons. To determine that, we examined the cell type specificity of expression for the FMRP bound mRNAs (Figure 5b). A large fraction (35%) of these mRNAs are in fact most abundant and most affected in neurons (cluster 1, Figure 5b-c, S3c-d). However, abundance does not seem to be critical for FMRP effect: a clear set of 34% of FMRP bound mRNAs are most abundant in non-neuronal cell types, but their downregulation is strongest in neurons (cluster 2, Figure 5b-c,S3c-d). The impact on FMRP bound gene expression may be further mediated by cell type specific expression of protein partners of FMRP ^5,76^, several of which are RNA binding proteins themselves. Many of these binding partners have functions in processes impacted in neurons or multiple cell types, such as splicing, cell signaling, vesicle transport and translation ^76^. We find no clear difference at RNA levels that suggests these genes contribute to the neuron-specific bound mRNA downregulation. Most FMRP binding partners are expressed at higher levels in neurons, oligodendrocytes and OPCs (Figure 5d). Taken together, this suggests that the neuron-specific effect likely results from the predominant expression of FMRP in neurons, previously shown by multiple immunostaining studies ^9,23,77,78^. We did not detect any difference in *Fmr1* expression across WT cells of different cell types (Figure S3b), which suggests the existence of a translational control mechanism behind FMRP levels in different cell types.

In conclusion, we find that FMRP bound mRNAs are preferentially downregulated in neurons, which are known to express higher levels of FMRP, and that this effect is also associated with the specificity of target mRNA expression.

## Neuron subtypes respond differently to the loss of FMRP

Many synaptic pathways are collectively impacted in *Fmr1*-KO neurons, as described above. We then examined if these and other pathways are uniformly or differently impacted in neuron subtypes. We focused on the three most abundant neuron subtypes: excitatory, interneurons and ganglionic eminence progenitors (Figure S1d). Indeed, we observed a common downregulation trend across most of the neuronal projection and synaptic pathways (Figure 6a).

**Figure 6.**
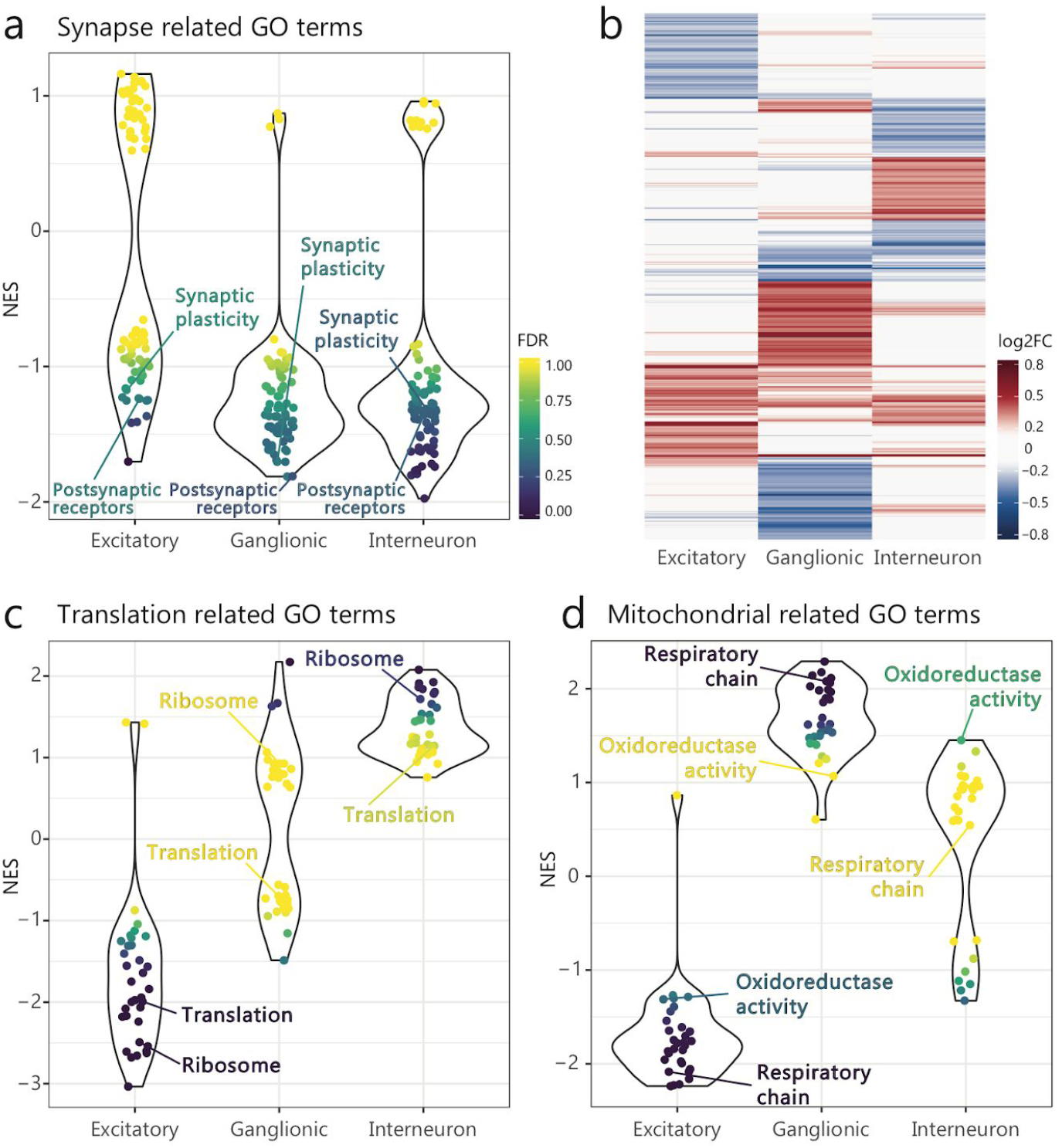
Neuron subtypes show diverse misregulated genes. **a)** Comparison of normalized enrichment scored (NES) across neuron subtypes for synaptic related GO terms (e.g. Synaptic plasticity = Regulation of synaptic plasticity, GO:0048167; Postsynaptic receptors = Regulation of postsynaptic membrane neurotransmitter receptor levels, GO:0099072). **b)** Overlap of top 5% upregulated (red) or downregulated (blue) genes in neuron subtypes. **c-d)** Comparison of normalized enrichment scores (NES) across neuron subtypes for translation related GO terms (c: e.g. Ribosome, GO:0005840; Translation, GO:0006412) or mitochondrial function related GO terms (d: e.g. Respiratory chain, GO:0070469; Oxidoreductase activity, GO:0016491).

However, we detected little overlap in the top dysregulated gene sets in these subtypes (Figure 6b). Firstly, we found that the increased expression of genes related to ribosomes and translation is unique to interneurons. In fact, excitatory neurons show the reverse trend, with downregulation of these genes (Figure 6c). Secondly, mitochondrial pathways are also affected in opposite ways in different neuronal subtypes. They are downregulated in excitatory neurons while upregulated in interneurons and ganglionic progenitors (Figure 6d). This reveals a critical difference in the effect of FMRP loss on different neuronal subtypes. It has been observed that mTOR signaling is increased in *Fmr1*-KO mice ^39,40^ as well as in humans with FXS ^79^. As mentioned previously, mTOR signaling results in increased metabolic activity and, in particular, in upregulation of translation activity (and ribosomal genesis) as well as in upregulation of mitochondrial processes ^37,38^. We observe such an increase only in inhibitory neurons, suggesting that they are the subtype affected by the hyperactivation of mTOR signaling. On the other hand, the downregulation of these pathways in excitatory neurons suggests that in this subtype there is a previously underappreciated dampening of mTOR activity.

Lastly, the top downregulated genes in interneurons are enriched for genes involved in pathways such as neuron maturation (GO:0042551, p<0.001, hypergeometric; e.g. App, Nrxn1) and synaptic vesicle localization (GO:0097479, p<0.01, hypergeometric; e.g. Adcy1, Apba1 and Syn2), which tend to be higher expressed at later stages of neuronal development ^80–83^. Conversely, top downregulated genes in excitatory neurons (other than ribosomal and mitochondrial related genes), include many regulators of early stage neurogenesis (e.g. Slc1a3, Vcan, Tpi1, Vim, Lrrc17, Sox9 and Ppap2b) ^84–88^. The contrasting natures of genes that are most downregulated in excitatory versus inhibitory neurons (neurogenesis vs neuron maturation), together with the opposite responses of the ribosomal and translational genes, which decrease with neuron differentiation ^89^, may reveal a misalignment of developmental progress between these two major neuronal subtypes. This misalignment may contribute to the excitatory-inhibitory imbalance proposed for FXS and other autism disorders ^90^.

## Discussion

Our study presents the first attempt to dissect the cell type specific contributions to FXS pathology using the power of single cell RNA-Seq, and reveals cell type specific alterations that could be masked in global measurements. We found that FMRP loss results in overall small changes to individual gene expression, however there is a broad misregulation of functionally related genes. We have described several novel findings resulting from the ability to dissect cell type specific effects in the FX mouse model including: i) Higher impact of FMRP loss in neurons; ii) Changes in abundance of cell-cell signalling genes that could increase environmental excitability; iii) Potential differences in mTOR activity across different cell types. We further discuss these findings below.

Neurons are markedly the most impacted cell type, and the processes associated with downregulated genes encompass critical functions for neuronal development, function and morphology. Neurons are also the only cell type where the set of mRNAs bound by FMRP in the mouse brain are distinctively downregulated. Recent evidence suggests that the degradation rates of FMRP bound mRNAs are higher in *Fmr1*-KO neurons, which results in their decreased abundance ^16^. The same post-transcriptional mechanism is likely behind the lower mRNA abundances detected in our single cell analysis and further strengthens the role of FMRP in the regulation of mRNA stability, possibly coupled to the regulation of translation rates ^91^.

A growing body of evidence supports the role of astrocytes in the pathology of neurological disorders ^92,93^. Astrocyte involvement in FXS in particular was previously demonstrated by co-culture studies where neurons exhibited decreased levels of synaptic proteins and abnormal dendritic morphology when grown with astrocytes from FXS mice ^94^. Our analysis of the *Fmr1*-KO single cell transcriptome reveals that many key receptors and secreted proteins are misregulated in astrocytes. These alterations suggest that *Fmr1*-KO astrocytes contribute to an environment of increased excitability, and could also impact neuronal development.

Multiple cell types display an upregulation of genes involved in translational and energy production processes, which are likely a downstream consequence of the increase in mTOR complex 1 signaling that has been previously detected in *Fmr1*-KO mice. The activation of this pathway is thought to be linked to the exaggerated mGluR5-LTD and a critical mechanism behind FXS pathology ^39,95^. Surprisingly, we find that cortical excitatory neurons in fact show an opposite downregulation trend for both types of processes, indicating that they respond to the loss of FMRP differently than inhibitory neurons and possibly are reflecting a decrease in mTOR activity. Indeed, stimulation of the N-methyl-D-aspartate receptor (NMDAR) decreases mTOR signaling activity ^96–98^. We find that NMDAR subunits are most highly expressed in excitatory neurons (Figure S4a). Further, we also find upregulation of one NMDAR subunit (Grin2b) specifically in *Fmr1*-KO excitatory neurons (Figure S4b, FDR=0.019). This increased expression of NMDAR in excitatory neurons, together with a glio and neurotransmitter environment that favors activation of NMDAR (Figure 3b-c), likely results in increased NMDAR signalling which suppresses mTOR activity and downstream pathways in excitatory neurons. On the other hand, inhibitory neurons exhibit increased metabolic gene expression which is concordant with mTOR hyperactivity. This implies that the current therapeutic strategies which propose components of the mTOR signaling pathway as targets for rescuing the cognitive and synaptic deficits in FXS have to consider that the outcome may be drastically different across neuron subtypes.

Finally, many of the neuronal downregulated pathways including neuron projection terms and vesicle transport were identified as downregulated both in RNA and protein levels by recent studies in human *in-vitro* neurons derived from embryonic or pluripotent stem cells ^99–101^. This suggests that the downstream molecular consequences of the absence of FMRP in neurons are well conserved and that the wealth of altered pathways we observed in the *Fmr1*-KO mouse model can provide a valuable resource for future studies in FXS patients and the design of therapeutic approaches.

## Methods

### Mice

Wild type (WT; FVB.129P2-*Pde6b*^*+*^ *Tyr*^*c-ch*^/AntJ; JAX stock 004828) and *Fmr1* knockout (*Fmr1*-KO; FVB.129P2-*Pde6b*^*+*^ *Tyr*^*c-ch*^ *Fmr1*^*tm1cgr*^/J; JAX stock 004624) mice at postnatal day 5 (P5) of both sexes were used. All experimental procedures followed the animal care protocol approved by the University of Massachusetts Medical School Institutional Animal Care and Use Committee (IACUC).

### Cortex dissection and dissociation

Three mice per genotype were used for single-cell collection. P5 mice were euthanized by decapitation and the brain was rapidly removed and placed in cold dissection buffer (1x HBSS). The brain was rapidly dissected to collect the cerebral cortex (both hemispheres) and the tissue was dissociated using the Papain dissociation system (Worthington Biochemical) following the manufacturer’s protocol with a 30 minute incubation.

### InDrop collection and scRNASeq library preparation

Dissociated cortical cells were resuspended in PBS, filtered through a 70 micron cell strainer followed by a 40 micron tip strainer, and counted with a Countess instrument (Life Technologies). The cells were further diluted to the final concentration (80,000 cells/mL) with OptiPrep (Sigma) and PBS, and the final concentration of OptiPrep was 15% vol/vol. A total of 2,000 to 5,000 cortical cells were collected per mouse and processed following the InDrop protocol ^29,102^. Final libraries containing ~1,200 cells were constructed and sequenced on an Illumina NextSeq 500 instrument, generating on average ~130 million reads per library (Table S3).

### Data processing

#### Cleanup, alignment and transcript quantification

Sequencing reads were processed as previously described ^103^ and the pipeline is available through GitHub (github.com/garber-lab/inDrop_Processing). Briefly, fastq files were generated with bcl2fastq using parameters *--use-bases-mask y58n*,y*,I*,y16n* --mask-short-adapter-reads 0 --minimum-trimmed-read-length 0 --barcode-mismatches 1*. Valid reads were extracted using the barcode information in R1 files and were aligned to the mm10 genome using TopHat v2.0.14 with default parameters and the reference transcriptome RefSeq v69. Alignment files were filtered to contain only reads from cell barcodes with at least 1,000 aligned reads and were submitted through ESAT (github.com/garber-lab/ESAT) for gene-level quantification of unique molecule identifiers (UMIs) with parameters *-wLen 100 -wOlap 50 -wExt 1000 -sigTest .01 -multimap ignore -scPrep*. Finally, we identify and correct UMIs that are likely a result of sequencing errors, by merging the UMIs observed only once that display hamming distance of 1 from a UMI detected by two or more aligned reads.

#### Dimensionality reduction and clustering

Gene expression matrices for all samples were loaded into R (V3.6.0) and merged. Barcodes representing empty droplets were removed by filtering for a minimum of 700 UMIs observed. The functions used in our custom analysis pipeline are available through an R package (github.com/garber-lab/SignallingSingleCell). Using the raw expression matrix, genes were selected for dimensionality reduction with a minimum expression of 3 UMIs in at least 1% of all cells. From those, the top 30% of genes with the highest coefficient of variation were selected. Dimensionality reduction was performed in two steps, first with a principal component analysis (PCA) using the most variable genes and the R package *irlba* v2.3.3 ^104^, then using the first 7 PCs (>90% of the variance explained) as input to a t-distributed stochastic neighbor embedding (tSNE; package *Rtsne* v0.15) with parameters *perplexity = 30, check_duplicates = F, pca = F* ^105^. Clusters were defined on the resulting tSNE 2-dimensional embedding, using the density peak algorithm ^106^ implemented in *densityClust* v0.3 and selecting the top cluster centers based on the *γ* value distribution (*γ* = *ϱ* × *δ; ϱ* = local density; *δ* = distance from points of higher density). Using known cell type markers, clusters that corresponded to erythrocytes or potential cell doublets were excluded and the remaining cluster were used to define 5 broad populations (Immune, Vascular, Astrocytes, Oligodendrocytes and Neuronal). Erythrocyte marker genes (*Hba-a1*, *Hba-a2*, *Hbb-b1*, *Hbb-bh1*, *Hbb-bh2*, *Hbb-bs*, *Hbb-bt*, *Hbb-y*, *Hbq1a*, *Hbq1b*) were excluded from the resulting expression matrix. Raw UMI counts were normalized separately for each batch of libraries (Table S3), using the function *computeSumFactors* from the package *scran* v1.12.1 ^107^, and parameter *min.mean* was set to select only the top 20% expressed genes to estimate size factors, using the 5 broad populations defined above as input to the parameter *clusters*. After normalization, only cells with size factors that differed from the mean by less than one order of magnitude were kept for further analysis (0.1×(***Σ***(*θ*/N)) > *θ* > 10×(***Σ***(*θ*/N)); *θ* = cell size factor; N = number of cells). The normalized expression matrix was used in a second round of dimensionality reduction and clustering. The top 20% variable genes were selected as described above, and a mutual nearest neighbor approach was used to correct for batch effects implemented in the *fastMNN* function of package *batchelor* v1.0.1 ^108^. To determine the number of dimensions used in the batch correction, the top 12 PCs that explained 90% of the variance were selected. The batch corrected embedding was used as input to tSNE, and the resulting 2D embedding was used to determine clusters as described above. Marker genes for each cluster were identified by a differential expression analysis between each cluster and all other cells, using *edgeR* ^109^, with size factors estimated by *scran* and including the batch information in the design model. Known cell type markers were used to determine 7 main populations (Immune, Endothelial, Pericytes, Ependymal, Astrocytes, Neuronal and Oligodendrocytes/OPCs). Each of these populations was independently re-clustered following the same procedure described above, to reveal more specific cell types and remove potential cell doublets (Figure S1). All R scripts for the steps described here are available through GitHub (github.com/elisadonnard/FXSinDrop).

#### Differential expression and GSEA analysis

Differentially expressed (DE) genes between genotypes were identified per cell type using *edgeR*, with size factors estimated by *scran* and including the batch information in the design model. Only genes that had at least one UMI in 15% of the cells of that type were used in the analysis. Genes were considered DE when they showed an FDR<0.01 and at least 1.15x fold change. Complete DE results can be found in Table S1.

The gene set enrichment analysis (GSEA v3.0)^110^ was performed using a ranked gene list constructed from the *edgeR* result. Genes were ranked by their reported fold change (logFC). GSEA was run via command line with parameters *--set_max 3000 -set_min 30*. The reference of GO term annotations for mouse Entrez GeneIDs was obtained from the *org.Mm.eg.db* package v3.8.2. GO terms were considered significantly altered if they showed an FDR<0.01. A total of 863 Gene Ontology (GO) terms had significant alteration (FDR<0.01) in one or more cell types. GO terms with high similarity were collapsed into 538 non-redundant categories (collapsedNAME, Table S2), using the function *plot_go_heatmap* from the package *SignallingSingleCell*, which compares the leading edge list of genes for each enriched pathway and creates a unified term if the overlap is greater than 80%. Collapsed categories were used to compare the results from different cell types (Figures 2a-b, S2c-d).

#### FMRP bound mRNA expression analysis

The normalized expression matrix was subset to select only WT cells and FMRP-bound mRNAs defined by Darnell *et al*. ^72^ expressed by at least one cell (n=743). Cells were ordered first based on cell type assignment and hierarchically clustered based on the expression of these mRNAs (*dist* function *method=”euclidian”*; *hclust* function *method=”average”*). Genes were clustered using *k-means* (k=7) followed by hierarchical clustering within the k-means cluster. Resulting clusters were merged based on the cell type pattern of expression to obtain the final three clusters (Neuronal, Non-neuronal, Non-specific).

---

**Figure S1.**
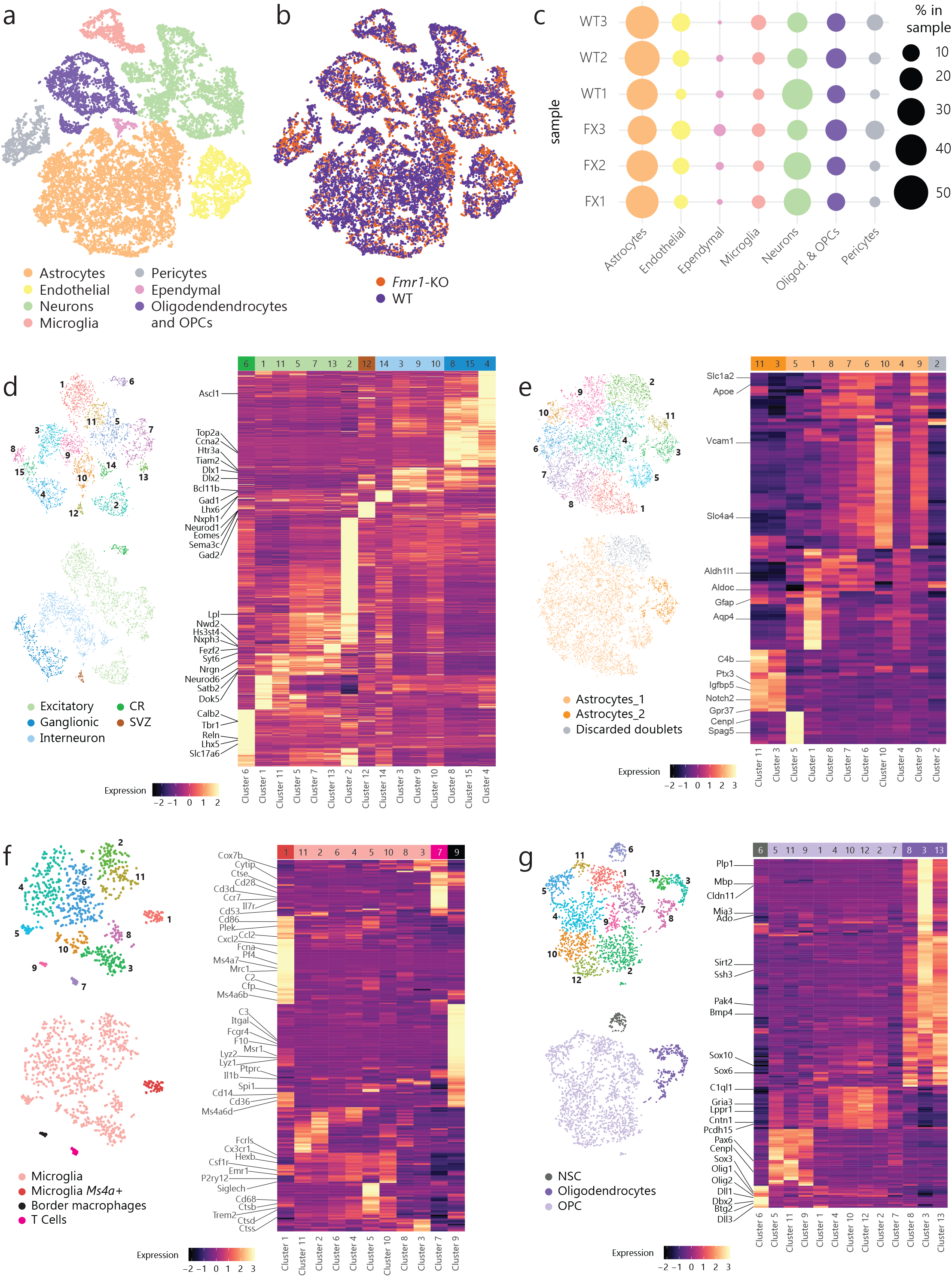
**a)** tSNE representation of cells colored by major cell type. **b)** tSNE representation of cells colored by genotype. **c)** Cell type composition of each collected sample. **d-g)** Unbiased sub-clustering results and subtype identification for each of the main cell types: **d)** Neurons; **e)** Astrocytes; **f)** Microglia and other immune cells; **g)** Oligodendrocytes and OPCs.

**Figure S2.**
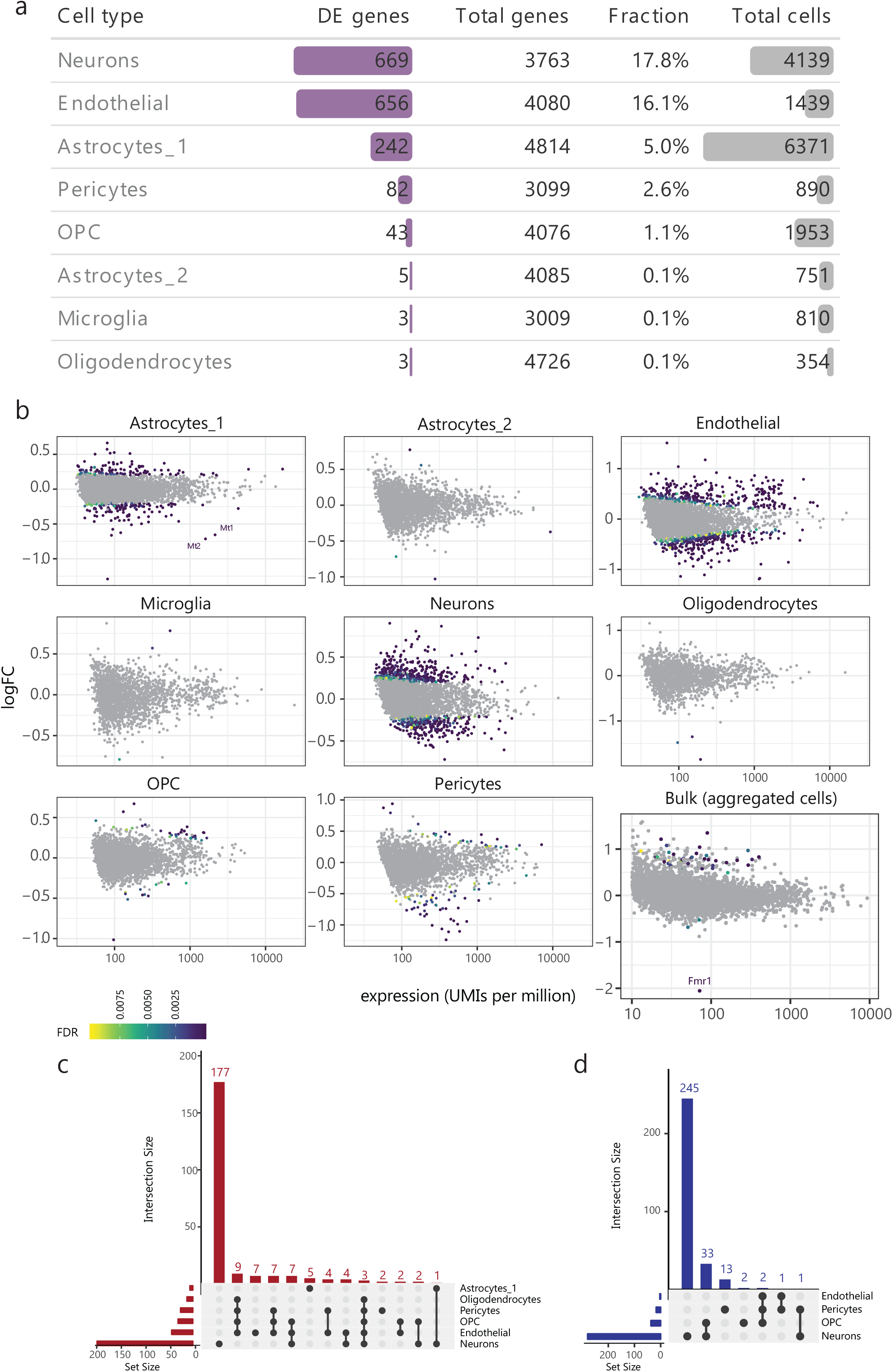

**a)** Differentially expressed (DE) genes identified per cell type. **b)** Plot of the log2 fold change (logFC) for each gene per normalized expression in aggregated WT cells (UMIs per million). **c-d)** UpSetR diagrams showing overlap between GO terms identified as significantly upregulated (c) or downregulated (d).

**Figure S3.**
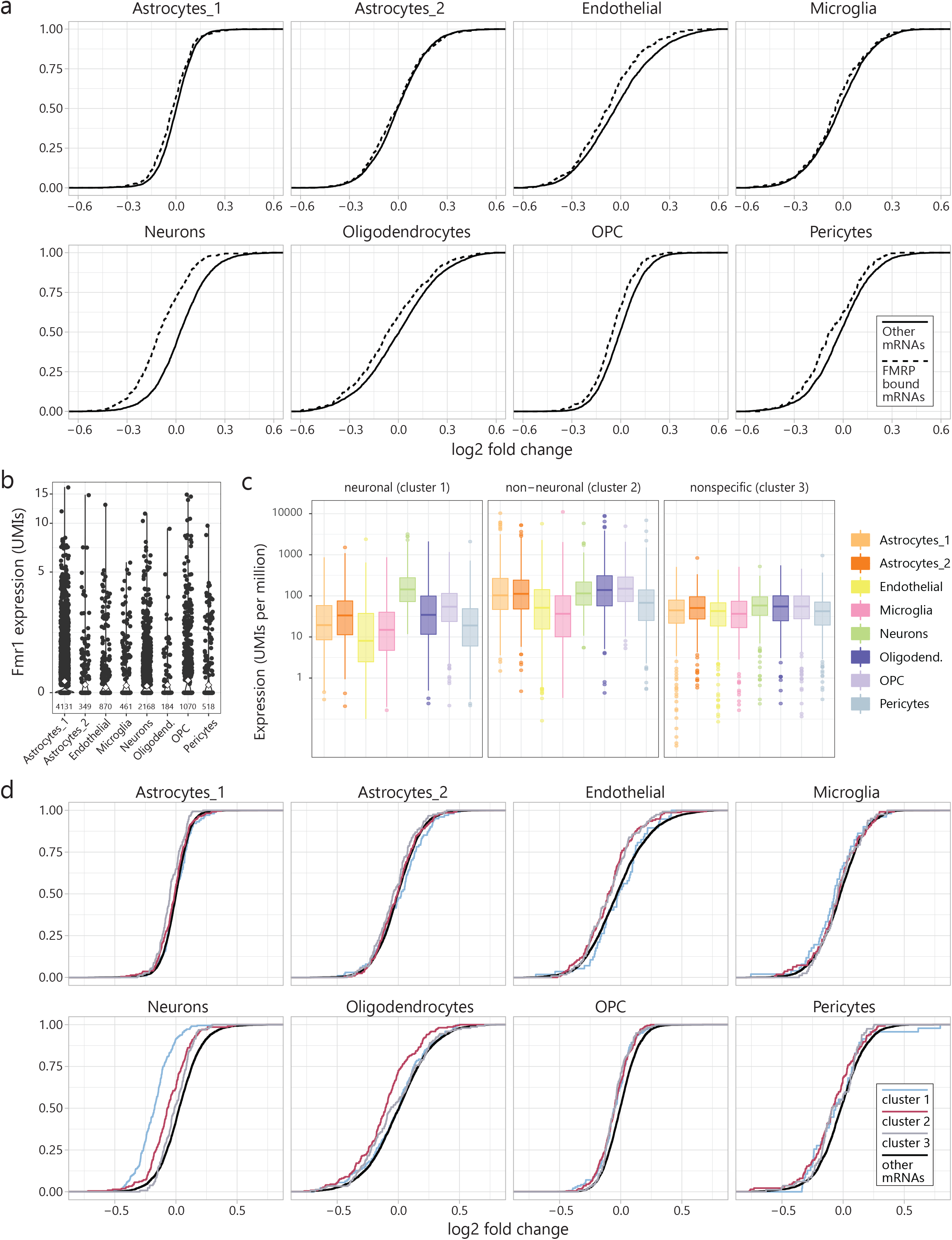
**a)** Cumulative distribution plot of the log2 fold change between *Fmr1*-KO and WT gene expression in each cell type. Dashed lines represent genes annotated as FMRP bound by Darnell et al. expressed in that cell type. Full lines represent all other expressed genes in that cell type. **b)** *Fmr1* expression (UMIs) in WT cells of each cell type. **c)** Aggregated expression (UMIs per million) in WT cells for each cell type of all FMRP bound genes that are classified as neuronal (cluster 1), non-neuronal (cluster 2), or non-specific (cluster 3) in terms of their pattern of expression (heatmap clusters, Figure 5b). **d)** Cumulative distribution plot of the log2 fold change between *Fmr1*-KO and WT gene expression in each cell type. Colored lines represent each specific subset of expressed genes genes: Neuronal FMRP bound mRNAs, which show highest expression in neurons (blue, cluster 1, Figure 5b); Non-neuronal FMRP bound mRNAs, which show highest expression in non-neuronal cell types (pink, cluster 2, Figure 5b); Non-specific FMRP bound mRNAs (grey, cluster 3, Figure 5b), which are expressed at similar levels by all cortical cell types; and all other mRNAs expressed in neurons (black).

**Figure S4.**
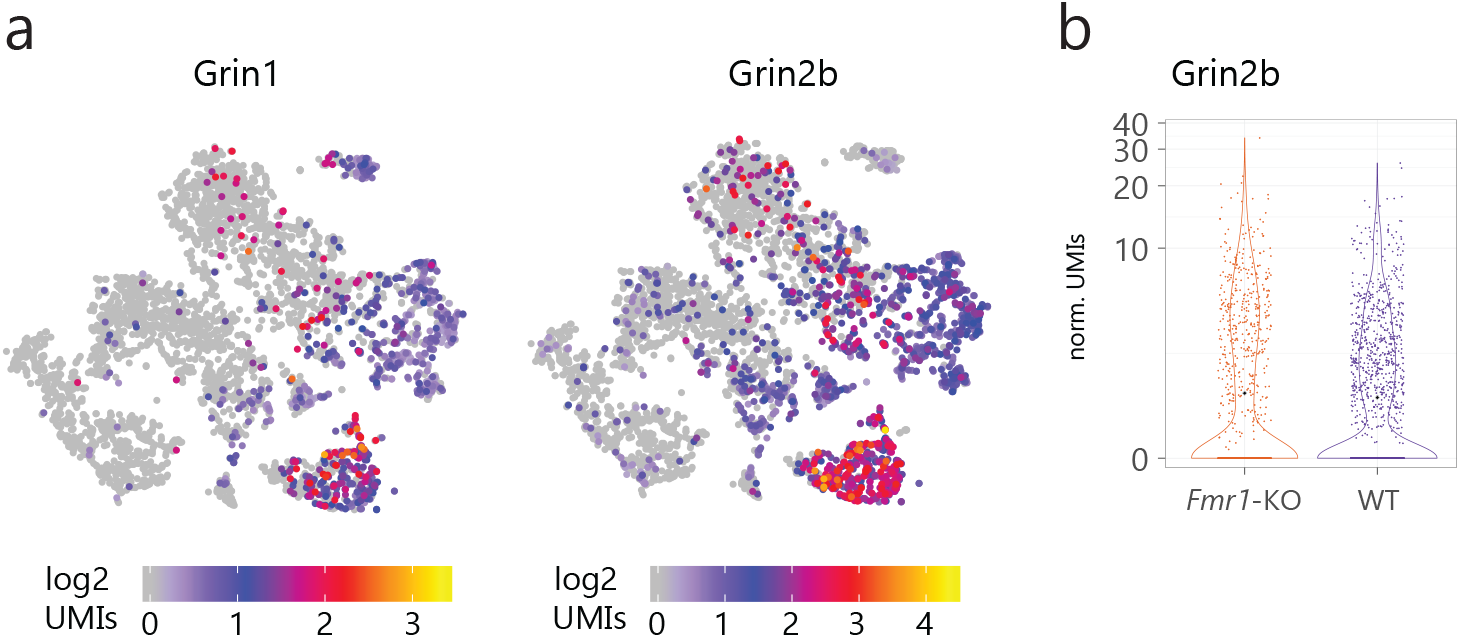
**a)** tSNE representation of neurons colored by expression level of NMDAR subunits detected, Grin1 (left) and Grin2b (right). **b)** Expression level (normalized UMIs) in *Fmr1*-KO or WT excitatory neurons for Grin2b.

